# Large-scale identification of microsatellite loci from multiple otter (Mammalia, Carnivora, Lutrinae) species using whole genome sequence data

**DOI:** 10.1101/2025.04.14.648774

**Authors:** Vera de Ferran, Channelle Ee Eun Chua, Philip Johns, Eduardo Eizirik, Klaus-Peter Koepfli

## Abstract

The development of molecular studies on elusive, rare, and/or poorly known species faces challenges due to the lack of suitable markers. Species-specific microsatellite markers minimize bias, offer better performance, and are cost-effective, aiding the development of population genetic studies. The use of whole-genome sequences allows for the development of species-specific microsatellite markers and their survey in closely related species, enabling the discovery of shared markers that can facilitate comparative studies. Lutrinae includes 14 extant species of otters. Despite their worrisome conservation status, due to inherent characteristics of these species that make their study difficult by traditional methods, many of them lack reliable population genetic data, limiting conservation efforts. In this study, we employed a multi-taxon approach to identify a large number of novel microsatellite loci for 11 of the 14 otter species, assessing whether the identified loci were shared among different taxa. We identified a total of 23,320 microsatellite loci across 11 species, which were reduced to 12,573 after stringent filtering criteria. Primer design was completed successfully for 420 and 259 unique loci, considering two minimum melting temperatures. We validated marker efficiency by testing the 81 loci designed for two Asian species. Of these, 51 loci yielded reliable microsatellite genotypes in both species, with 34 showing allelic variation in at least one of them. These results demonstrate that these markers are applicable in empirical genotyping for both their target and closely related ones.

## INTRODUCTION

Molecular approaches have revolutionized the fields of genetics, ecology, and evolutionary biology, and are routinely applied in surveys, monitoring programs and status assessments of threatened organisms (Schwartz et al. 2007). However, the application of such methods to wildlife species that are rare, elusive, and/or poorly studied is often hindered by the paucity of appropriate molecular markers for efficient genetic analysis. Therefore, developing molecular tools that are specifically designed for poorly known taxa is a useful avenue to enable genetic, ecological, and conservation-oriented studies targeting these species.

Of the various molecular markers developed in the last four decades to survey natural populations, microsatellite loci have been shown to be particularly informative for multiple types of problems (Frankham et al. 2002). For example, they have been used in many different taxa to assess levels of genetic diversity, population structure and demographic connectivity, as well as to perform individual-based ecological, behavioral, and forensic analyses (e.g., Trinca et al. 2013, Miller et al. 2014, Ishida et al. 2018, Harper et al. 2018, Sun et al. 2020). Despite their usefulness, the routine application of microsatellites to address such problems in wildlife species has been historically hampered by the lack of markers developed specifically for the target organism, often requiring their transfer (i.e., use of the same primers with adjusted PCR conditions) from better-known, related species (Frankham et al. 2002). Although this approach is often successful, species-specific markers are expected to show better performance in terms of variability (informative content) and PCR efficiency (which allows, for example, improved recovery of molecular data from degraded field-collected materials such as feces and hairs). In addition, their use should minimize ascertainment bias associated with using heterologous loci (Li and Kimmel 2013).

Despite the increasing application of reduced representation and low-coverage whole genome sequencing approaches to analyze single nucleotide polymorphisms (Fuentes-Pardo and Ruzzante 2017), microsatellites remain a cost-effective and viable tool for population genetic analyses of animal and plant populations, especially in species with reduced genetic diversity (Hauser et al. 2021) or that are being genetically surveyed for the first time. Genome sequencing has greatly facilitated the development of species-specific microsatellite markers by enabling the rapid computational discovery of repetitive sequences from genomic data (Benson 1999). Although this strategy has now been used for many systems (e.g., Kumari et al. 2019, Liu et al. 2019, Latorre-Cardenas et al. 2020), and it is starting to become more common with the increase in the availability of genomic resources, it is still not commonly applied to whole-genome sequences from multiple species of the same clade (but see Liu et al. 2017, Xu et al. 2018, Bhat et al. 2018, Manee et al. 2020 for examples). Such an approach would allow the parallel survey of microsatellites in closely related species, enabling the discovery of shared markers that can facilitate comparative studies.

An interesting clade in which this approach can be assessed is the mustelid subfamily Lutrinae, which comprises 14 extant species of otters. The current IUCN Red List of Threatened Species (IUCN 2022) categorizes otter species as: ‘Least Concern’ (*Lontra canadensis*, North America); ‘Near Threatened’ (*Aonyx capensis*, Sub-Saharan Africa; *Aonyx congicus*, Central Africa; *Hydrictis maculicollis*, Sub-Saharan Africa; *Lontra longicaudis*, South America; *Lutra lutra*, Eurasia and North Africa); ‘Vulnerable’ (*Aonyx cinereus, Lutrogale perspicillata*, South and Southeast Asia); and ‘Endangered’ (*Enhydra lutris*, North Pacific Rim; *Lontra felina*, Pacific coast of South America; *Lontra provocax*, southern South America; *Lutra sumatrana*, Southeast Asia; *Pteronura brasiliensis*, South America). The recently recognized *Lontra annectens* of Central America (de Ferran et al. 2024) has not been assessed yet by the IUCN. Despite their worrisome conservation status and the estimated reduction in census sizes for most of these species, there is no reliable data on demographic, ecological and genetic aspects for most of these taxa throughout most of their respective geographic distributions (*Enhydra lutris* being a notable exception, e.g., Larson et al. 2021, Beichman et al. 2023), which limits conservation planning and action.

Otters have several characteristics that make their study difficult by traditional methods: most species (except *Pteronura brasiliensis* and *Hydrictis maculicollis*) do not present a coat pattern (e.g. spots or stripes) that allow individual identification; they are semi-aquatic, limiting the use of camera-traps or radio-telemetry methods and also making it difficult to capture and immobilize these animals; and several species are secretive and not commonly observed in the wild, in addition to being crepuscular/nocturnal. For these reasons, most studies on these species are based on indirect records, such as using feces and latrines as sources of information on presence, diet, home range size, and activity (e.g., Crowley et al. 2012; Rivera et al. 2019). However, these methods are quite limited for population studies due to the impossibility of identifying individuals with morphology-based methods, along with the difficulty in differentiating species in areas where more than one otter species occur in sympatry (e.g., Southeast Asia; Koepfli et al. 2008).

One way to overcome these limitations is to employ molecular methods that provide powerful sources of species-specific as well as population- and individual-level information from various types of biological samples. In this context, the recent availability of whole-genome sequences from 13 of the 14 species in Lutrinae (de Ferran et al. 2022) has opened the possibility of performing surveys of microsatellite loci across this clade. In this study, we have employed this multi-taxon approach to identify novel microsatellite loci for 11 of the 14 otter species. We validated their efficiency by testing them on two species, providing a valuable resource that should help enhance genetic and ecological studies targeting these organisms.

## MATERIAL AND METHODS

### Genomic dataset

To conduct a large-scale, standardized survey of microsatellite loci in the subfamily Lutrinae, we analyzed whole-genome sequences from 13 extant otter species. For 11 species (*Aonyx cinereus, Aonyx capensis, Aonyx congicus, Hydrictis maculicollis, Lontra canadensis, Lontra felina, Lontra longicaudis, Lontra provocax, Lutra lutra, Lutra sumatrana* and *Lutrogale perspicillata*), we used genomes that were previously reported by our group (de Ferran et al. 2022), while for *Pteronura brasiliensis* and *Enhydra lutris*, we used publicly available genome assemblies (Beichman et al. 2019). For two species (*Lutra sumatrana* and *Aonyx congicus*), the genome sequences were generated from museum specimens, and therefore had much lower depth of coverage and more missing data than the other species (see de Ferran et al. 2022). All 13 genomes were mapped against the Eurasian otter (*Lutra lutra*) reference genome (Mead et al. 2020) following the protocol reported by de Ferran et al. (2022). We generated consensus sequences of all genomes using ANGSD 0.921 (Korneliussen et al. 2014) with the parameters doFasta= 2, doCounts= 1, explode= 1, setMinDepth= 10 and minMapQ= 20.

### Microsatellite mining and filtering

To identify tandem repeats in each species’ genome, we used the Tandem Repeats Finder program (Benson 1999) with default parameters: matching weight = 2, mismatching penalty = 7, indel penalty = 7, match probability = 80, indel probability = 10, minimum alignment score to report = 50, maximum period size to report = 4, to include flanking sequence (-f) and data file (-d), and maximum TR length expected in millions (-l) = 2. Considering the minimum score and the matching weight, the minimum number of required units in the tandem array was seven. We filtered the results stringently by keeping only tetranucleotides with complete and perfect repeats, and without any missing data in the target sequence or in either of the flanking regions (50 bases on each side of the repetitive array). Tetranucleotide loci were chosen due to their reduced stutter artifacts and simpler allele size determination (e.g., Edwards et al. 1991; Guichoux et al. 2011), despite potentially lower mutation rates compared to loci with smaller repeat motifs (e.g., dinucleotides; Chakraborty et al. 1997).

The repeat regions that were retained after these filtering steps were then assessed in terms of the efficiency of PCR primer design targeting their flanking sequences. Our aim was to develop microsatellite loci for each species with minimal need for PCR optimization by multiple end users (Robertson and Walsh-Weller 1998). This was conducted with Primer3 (Koressaar and Remm 2007, Untergasser et al. 2012) using the following parameters:

PRIMER_TASK=generic, PRIMER_PICK_LEFT_PRIMER=1, PRIMER_PICK_INTERNAL_OLIGO=0, PRIMER_PICK_RIGHT_PRIMER=1, PRIMER_OPT_SIZE=20, PRIMER_MAX_SIZE=20, PRIMER_NUM_RETURN=1, PRIMER_OPT_TM=60.0, PRIMER_MAX_TM=60.1, PRIMER_MIN_TM=59.9, PRIMER_PRODUCT_SIZE_RANGE=80-400. To assess the effect of such a stringent range of melting temperatures (TM), we performed a second round of analysis with the same parameters, changing only PRIMER_MIN_TM to 59.8. We chose these temperature thresholds because of the increased specificity of PCR during primer annealing at higher temperatures (Roux 1995, Hecker and Roux 1996).

In addition to discovering microsatellite repeats in each species, we also assessed whether the identified loci were shared among different otter species. To do this, we compared the genomic coordinates of the identified repeats, since all species had their genome sequence consensus constructed using the same Eurasian otter reference genome assembly. We also compared the repeat motif and the flanking sequences of each shared locus to assess their variation across species.

### Empirical validation of microsatellite genotyping

To ascertain that microsatellites identified with our approach could be reliably genotyped, we empirically tested the markers developed for two Asian species, the smooth-coated otter (*Lutrogale perspicillata*) and the small-clawed otter (*Aonyx cinereus*) for which samples had been collected and were available to be used for this study. We optimized the 59.9°C set of primers designed for each of these species using genomic DNA from only one individual of each, isolated previously with Qiagen DNeasy Blood & Tissue DNA isolation kits (www.qiagen.com) from tissue samples obtained from the cryo-collection of the Lee Kong Chian Natural History Museum, Singapore (lkcnhm.nus.edu.sg; *Lutrogale perspicillata* sample LKCNHM 119390 and *Aonyx cinereus* sample LKCNHM 118524). Additionally, we tested a set of 27 primers (7 from *Aonyx cinereus*, 11 from *Lutrogale perspicillata*, and 9 duplicated, occurring in both species) designed based on a random set of imperfect repeats, to assess whether they can also yield useful markers (Table S5).

All PCRs were conducted in 20 µL reaction volumes, using 10 µL Promega 2x Taq master mix (www.promega.com), 2 µL DNA eluate, and 1.0 µl each of 10 µM forward and reverse primers, on a Bio-Rad T100 thermocycler (www.bio-rad.com). First, we conducted a touchdown PCR with the following thermal cycling profile: 94ºC for 180s as an initial denaturation step, followed by 30 cycles of 30s at 94ºC for denaturation, 30s of annealing temperatures (T_a_) starting at 65°C and dropping 0.5°C per cycle, and 30s at 72ºC for extension, followed by 20 similar cycles with T_a_ of 50ºC, then 240s at 72ºC, and 4ºC forever. PCR products were run on a 1% agarose gel to assess quality and were classified as either no product, multiple banding pattern indicative of non-specific binding, or a clear single-band PCR product (N, M, or Y, respectively). If we obtained a clear PCR product, or if a multiple banding pattern was not severe, we conducted gradient PCR to determine the optimum annealing temperature, with 30 cycles of 30s each of 94ºC, T_a_, and 72ºC, where the gradient annealing temperatures ranged from 65° to 50°C. We conducted the final PCRs using similar thermal cycling conditions to the gradient reaction conditions, at the optimized annealing temperature for each primer pair, and using forward primers labelled with fluorophores 6-FAM, HEX or TET. These PCR products were submitted to 1st-Base Asia, Singapore (www.base-asia.com) where they were run on an Applied Biosystems capillary sequencer (https://www.thermofisher.com). Microsatellite allele sizes were scored using the Microsatellite v1.4.7 plugin in the Geneious Prime software (https://www.geneious.com).

## RESULTS AND DISCUSSION

Using enrichment, cloning, and Sanger sequencing of genomic libraries, limited numbers of microsatellite loci have been previously characterized in several otter species including *Lutra lutra* (Dallas and Piertney, 1998; Huang et al. 2005), *Enhydra lutris* (Kretschmer et al. 2009), *Lontra canadensis* (Beheler et al. 2005) and *Pteronura brasiliensis* (Ribas et al. 2011). Microsatellite loci are abundant in eukaryotic genomes (Chanzi and Georgakopoulos-Soares, 2024). These loci can be efficiently identified from genome assemblies or sequencing data using bioinformatics tools like Tandem Repeats Finder (Benson, 1999). In this study, we identified a total of 23,321 tetranucleotide microsatellite loci within the genomes of 12 of the 13 otter species we analyzed (all but *A. congicus*). This total was obtained by summing the identified loci in each species, and therefore it includes markers that were duplicated due to being found in more than one species (Table S1). Among these loci, due to the enforcement of stringent filtering criteria, only 15 were shared among the 11 species with higher depth of coverage (i.e., excluding the genomes generated from museum samples, Table 1, Table S2). Of the two otters represented by museum samples, only one locus was identified for *L. sumatrana*, and none for *A. congicus*, indicating that the search criteria that we enforced were too stringent for lower-coverage genomes with larger amounts of missing data. However, it is noteworthy that each of them has a sister-species (*L. lutra* and *A. capensis*, respectively) that yielded a large amount of identified loci, which are likely to be applicable on samples obtained from *L. sumatrana* and *A. congicus*.

**Table 1.** Total number of identified microsatellite loci per species for the 11 species with higher-coverage genomes, and the subsets for which PCR primers could be successfully designed with two different settings for the minimum melting temperature, Tm (expressed in °C). The single locus identified for *L. sumatrana* (see text) is not shown (see Tables S1, S2).

| Species | Total | Only perfect repeats | Successful primer design. Minimum T<sub>m</sub>=59.8°C | Successful primer design. Minimum T<sub>m</sub>=59.9°C |
| --- | --- | --- | --- | --- |
| *Aonyx cinereus* | 2,157 | 1,136 | 56 | 28 |
| *Aonyx capensis* | 2,232 | 1,133 | 67 | 42 |
| *Enhydra lutris* | 2,436 | 1,332 | 85 | 47 |
| *Hydrictis maculicollis* | 1,590 | 794 | 44 | 27 |
| *Lontra canadensis* | 1,556 | 854 | 39 | 28 |
| *Lontra felina* | 1,182 | 610 | 32 | 16 |
| *Lontra longicaudis* | 2,079 | 1,180 | 83 | 54 |
| *Lontra provocax* | 1,027 | 510 | 23 | 12 |
| *Lutra lutra* | 5,359 | 3,080 | 179 | 105 |
| *Lutrogale perspicillata* | 2,308 | 1,213 | 66 | 43 |
| *Pteronura brasiliensis* | 1,394 | 731 | 57 | 31 |
| **Total** | **23,320** | **12,573** | **731** | **433** |

Because some of the loci presented occasional differences in the repeat motif, which could be exclusive to one species or occur in multiple species (Figure 1), we applied another filter layer to exclude these cases, leaving only perfect repeat arrays to be considered in the downstream step of primer design. Still, because these variations may be of interest to researchers working on these species, these data are also included in the Supplementary Material. After this new filter, there were 12,573 sequences left (Table 1; Figure 2; Table S2), with no locus left for *L. sumatrana*, and no shared loci among all the 11 remaining species (Table S2). At the same time, if a smaller set of species was assessed (e.g., those occurring on the same continent), some overlap in the retrieved loci could be discerned, opening the possibility of designing markers that can be applied to sets of sympatric otter species.

**Figure 1.**
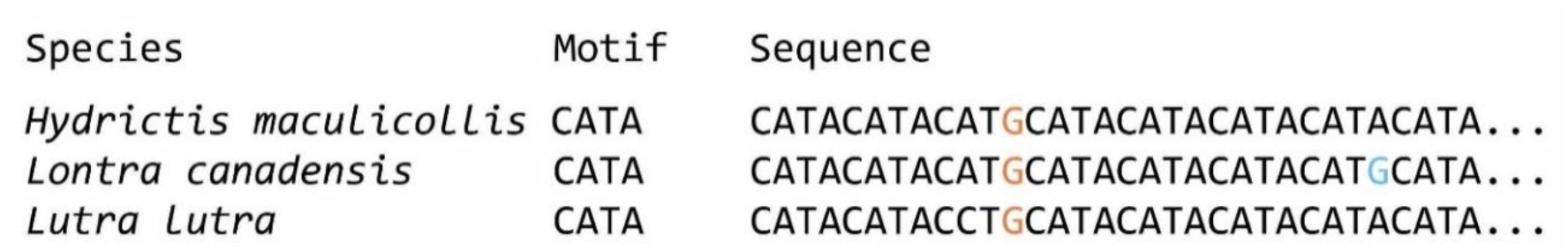
Example of filtered differences in relation to the repeat pattern present in several species (orange) or exclusive to one (blue).

**Figure 2.**
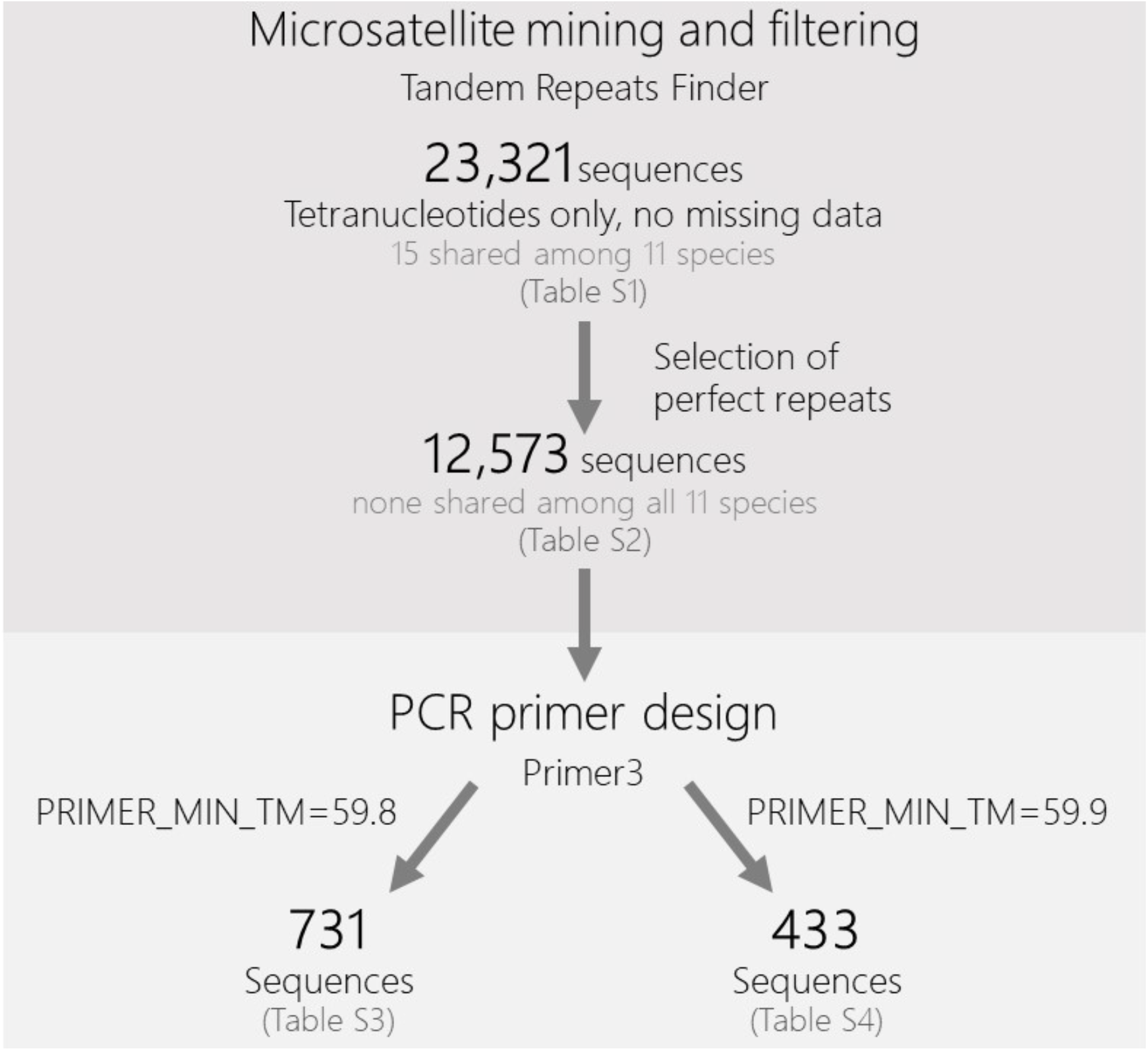
Flow chart depicting the sequential steps conducted in this study, from the identification of microsatellite loci to the filtering of perfect repeats and the design of PCR primers for these markers.

The primer design step was completed successfully for 731 sequences (representing 420 unique loci) for the 59.8°C minimum melting temperature (Table S3) and 433 sequences (representing 259 unique loci) for the 59.9°C minimum melting temperature (Table 1, Table S4). This substantial difference in the number of retrieved loci with only a change of 0.1°C in the minimum Tm illustrates the impact of this parameter when performing such genome-wide surveys of microsatellite markers and highlights the need to consider it carefully. Overall, the 59.9°C minimum melting temperature group presented 57 repeat patterns, and the 59.8°C one presented 68 patterns. For both, the most common repeat patterns were the ones containing AT combinations (AAAT, ATTT, TAAA, and TTTA).

Although we used a different *Lutra lutra* individual relative to the one employed for the reference genome (to minimize the potential influence of the mapping reference), *Lutra lutra* was the species with the largest number of identified loci. The reference seemed to influence the number of retrieved loci per species, with species closer to the reference (i.e., with a more recent divergence) having, overall, a larger number of identified loci and a lower amount of missing data (hence, fewer sequences were excluded during the filtering steps). This in turn led to the observed bias in the success rate for the reference species (see Table 1). In spite of this bias, we still obtained a substantial number of microsatellite loci for every surveyed species (>20 loci with primers designed with a minimum Tm=59.9°C; >40 loci with a minimum Tm=59.8°C), demonstrating the potential of this approach to identify such sequences across a whole group of related species, while enforcing strict criteria that maximize the species-specificity and informative potential of these markers.

We assessed the number of shared loci among species from the same continent, considering only markers for which PCR primer design was successful (Figure 3). The observed trends were similar for both assessed melting temperatures, with several combinations of shared markers among potentially sympatric species. For example, in Eurasia, there were 13 shared loci among the three surveyed species when the minimum Tm was 59.9°C, and 20 shared loci with minimum Tm=59.8°C. In South America, where four species were compared, different combinations of shared loci could be retrieved. Among these, the two sympatric tropical species *Pteronura brasiliensis* and *Lontra longicaudis* shared 10 loci with minimum Tm=59.8°C, exemplifying a marker set that can be useful to investigate both species in the field.

**Figure 3.**
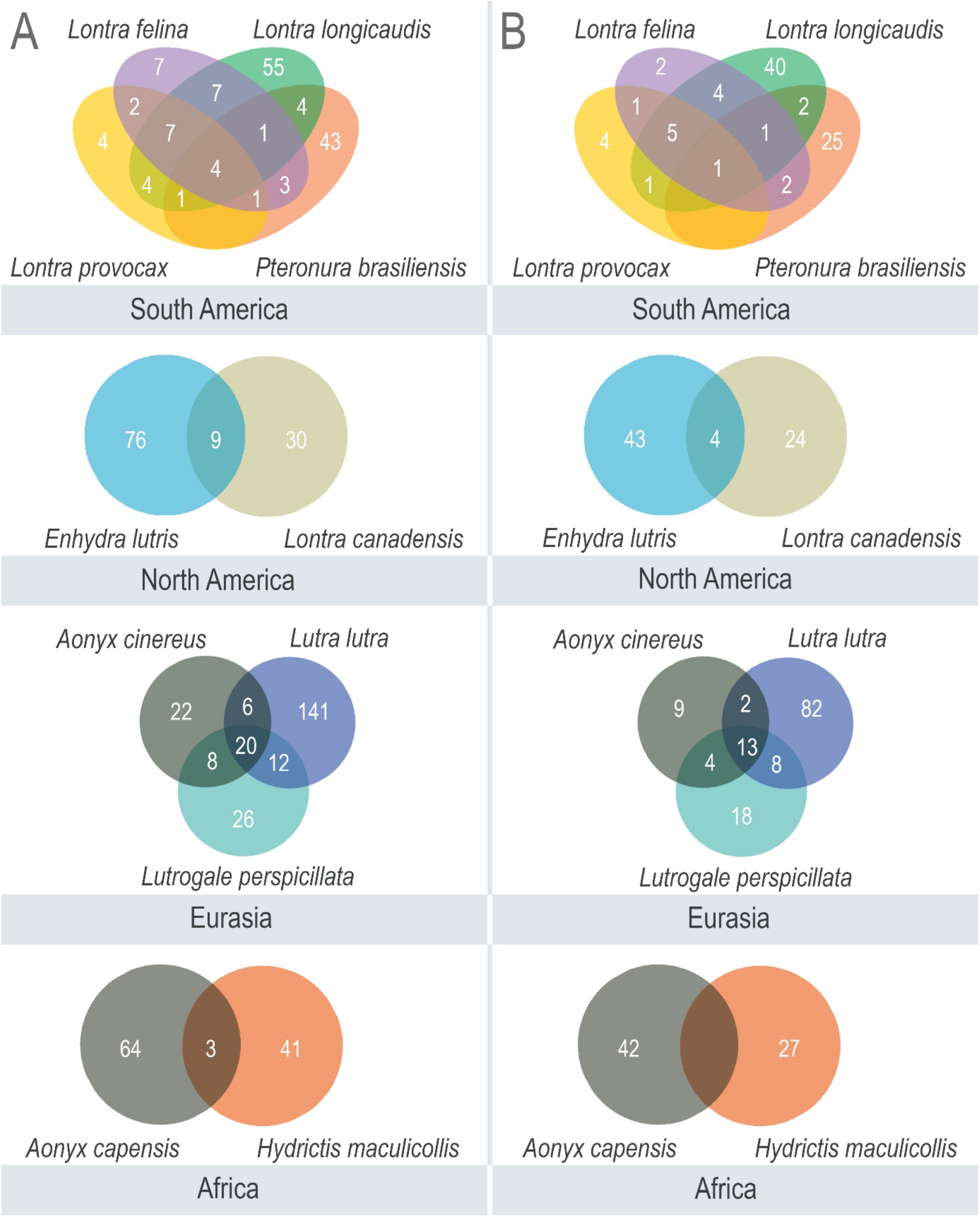
Venn diagrams depicting the number of microsatellite loci retrieved for otter species from different continents (including those that were shared among species), for which PCR primers could be successfully designed. A) Results obtained when the primer melting temperature (Tm) was restricted between 59.8°C and 60.1°C; B) Results obtained when the Tm was restricted to be between 59.9°C and 60.1°C.

Finally, the marker set with minimum Tm of 59.9°C identified for *Aonyx cinereus* (44 in total) and *Lutrogale perspicillata* (63 in total), based on perfect and imperfect repeats (Table S4, S5), was empirically tested in one individual of each of these species. Considering that these species had primers in common (Figure 3), the total number of loci/primer sets tested was 81 (18 designed for *A. cinereus*, 37 designed for *L. perspicillata*, and 26 common to both). Of these, 51 yielded successful microsatellite genotypes in both species (Table S6), and in 34 of them we observed heterozygosity in at least one of the otters (Table S6). Interestingly, of the nine loci designed exclusively for *A. cinereus* that showed variability, two were heterozygous in the target species and homozygous in *L. perspicillata*, while four showed the opposite pattern and three were heterozygous in both species. Similarly, of the 12 loci designed exclusively for *L. perspicillata* that showed variability, four were heterozygous in the target species and homozygous in *A. cinereus*, while five showed the opposite pattern and three were heterozygous in both species. We did not find any evident difference regarding the allelic size range between heterologous and homologous loci. These results demonstrate that these markers can be successfully applied in empirical genotyping, and that they are informative (i.e., variable) not only in their respective target species but also in another, closely related otter. However, we acknowledge that comparing monomorphic or polymorphic loci across multiple individuals of the two species would be more informative than comparing a single homozygous or heterozygous individual between these species.

Similar sets can be constructed for other combinations of otter species that occur in a given area, not only using the Tm range that we employed here, but also considering a broader suite of PCR parameters, starting from our complete list of identified tetranucleotide markers (Tables S1, S2). In addition, microsatellite marker sets can be designed to allow for multiplexing, thereby making data collection more efficient (Puckett 2017). Furthermore, when paired with markers containing species-specific diagnostic sites (e.g., Koepfli et al. 2008), our panel of microsatellite loci will empower genetic and population monitoring of otter species with overlapping distributions. The microsatellite loci we report in this study will need to be empirically assessed for polymorphism and information content by individual researchers and if they are applied to non-invasive samples such as hair and spraints, further optimization and testing may be required. We believe this resource should be useful to facilitate global research on otter ecology and genetics, accelerating the collection of field data with relevant implications for conservation planning on behalf of these species.

## ACKNOWLEDGMENTS

We thank Foo Maosheng (Lee Kong Chian Natural History Museum, Singapore) for access to samples. Financial support for this study was provided by CNPq/Brazil (grants 141172/2017-7 [awarded to V.d.F.] and 309068/2019-3 [awarded to E.E.]), the PUCRS/CAPES-PrInt Program (fellowship 88887.370464/2019-00 awarded to V.d.F.), the Ministry of Education through the Yale-NUS College start-up grant R-607-265-226-121 (awarded to C.E.E.C. and P.J.) and by the Yale-NUS College Centre for International & Professional Experience (awarded to C.E.E.C.). This study is a contribution of the National Institutes for Science and Technology (INCT) in Ecology, Evolution and Biodiversity Conservation, supported by MCTIC/CNPq/Brazil (proc. 465610/2014-5) and FAPEG/Brazil (proc. 201810267000023). We thank two anonymous reviewers who provided constructive comments on an earlier version of this manuscript.

